# Cannabinoid tetrad effects of oral Δ9-tetrahydrocannabinol (THC) and cannabidiol (CBD) in male and female rats: sex, dose-effects and time course evaluations

**DOI:** 10.1101/2021.05.11.443601

**Authors:** Catherine F. Moore, Elise M. Weerts

## Abstract

*Rationale* The legalization of medicinal use of *Cannabis sativa* in most US states and the removal of hemp from the Drug Enforcement Agency (DEA) controlled substances act has resulted in a proliferation of products containing Δ9-tetrahydrocannabinol (THC) and cannabidiol (CBD) for oral consumption (e.g., edibles, oils and tinctures) that are being used for recreational and medicinal purposes. *Objective* This study examined the effects of cannabinoids THC and CBD when administered orally on measures of pain sensitivity, body temperature, locomotor activity, and catalepsy (i.e., cannabinoid tetrad) in male and female Sprague Dawley rats. *Methods* Rats (N=24, 6 per sex/drug group) were administered THC (1-20 mg/kg), CBD (3-30 mg/kg), or sesame oil via oral gavage. Thermal and mechanical pain sensitivity (tail flick assay, von Frey test), rectal measurements for body temperature, locomotor activity, and the bar-test of catalepsy were completed. A separate group of rats (N=8/4 per sex) were administered morphine (5-20 mg/kg; intraperitoneal, IP) and evaluated for pain sensitivity as a positive control. *Results* We observed classic tetrad effects of antinociception, hypothermia, hyper- and hypolocomotion, and catalepsy after oral administration of THC that were long lasting (>7 hours). CBD modestly increased mechanical pain sensitivity and produced sex-dependent effects on body temperature and locomotor activity. *Conclusions* Oral THC and CBD produced long lasting effects, that differed in magnitude and time course when compared with other routes of administration. Examination of cannabinoid effects administered via different routes of administration, species, and in both males and females is critical to enhance translation.

## Introduction

Changes in federal regulation of hemp, and legalization of medicinal use of *Cannabis sativa* and its constituents in the majority of US states has resulted in a proliferation of cannabis products. Oral formulations of the cannabinoids Δ9-tetrahydrocannabinol (THC) and cannabidiol (CBD) are common for people using cannabis medicinally (e.g. Dronabinol/Marinol™, Epidiolex) and oral cannabis-derived products are widely marketed (e.g., edibles, oils and tinctures)(Spindle et al., 2019).

Preclinical research into cannabinoid effects has been ongoing for decades, with a majority of research to date employing injection methods (e.g., intraperitoneal, IP; subcutaneous, SC; intravenous, IV) for drug administration. Some of the foundational preclinical research into cannabinoids came from the development of the classic tetrad test, where behavioral effects of THC and synthetic cannabinoid receptor 1 (CB1R) agonists were linked with their pharmacological activity on CB1R. The test includes measures of pain sensitivity, body temperature, locomotor activity, and catalepsy. IP, IV, and SC administration of THC and other CB1R agonists produce phenotypic responses of antinociception, hypothermia, hypolocomotion, and catalepsy, and these effects are blocked with CB1 antagonists (Compton et al., 1993; Martin et al., 1991; Metna-Laurent et al., 2017). Preclinical evaluations of cannabinoid effects in the context of oral administration methods are limited. Furthermore, to date, studies have been conducted primarily in mice with behavioral assays conducted at few discrete timepoints post administration. To our knowledge, no studies have yet examined the time course of tetrad effects following oral administration of THC in male and female rats. As animal models are increasingly using oral routes of administration for cannabinoid exposure, data on the classic behavioral tetrad provide the basis for selection of pretreatment times and optimal dosing to assess behavioral outcomes.

Use of an oral route of administration is important from a translational approach. While IP injection and oral administration both undergo first pass hepatic metabolism, there are pharmacokinetic differences. When taken orally, THC is slowly and erratically absorbed (Grotenhermen, 2003; Newmeyer et al., 2017; Wall et al., 1983). A pharmacokinetic study of oral, subcutaneous injection, and vapor administration in rats found that oral administration resulted in long-lasting levels of THC in serum and brain, and the highest brain levels of THC compared to other routes of administration (Hlozek et al., 2017). The pharmacokinetic parameters of oral consumption of THC are further affected by the formulation (solution vs. capsule) and food (high fat food vs. fasted state). Further, cannabinoid receptors are distributed throughout the gastrointestinal tract and some cannabinoid effects may be mediated peripherally.

Therefore, this study sought to characterize the tetrad effects of THC (0-20 mg/kg) administered orally in a high fat vehicle in male and females rats over multiple hours post-administration. We also assessed the effects of oral CBD (0-30 mg/kg) on three of the four tetrad behaviors (antinociception, hyperthermia, locomotor activity). Cataleptic behavior was not assessed in animals treated with CBD, as prior studies demonstrated that CBD does not induce catalepsy, but instead has anticataleptic effects (Gomes et al., 2013).

## Materials and Methods

### Subjects

Adult male and female Sprague Dawley rats (Charles River, Wilmington, MA) were single housed in wire-topped, plastic cages (27 × 48 × 20 cm) with standard enrichment. The vivarium was on a 12hr reverse light/dark cycle (lights off at 9:00 a.m.) and was humidity and temperature controlled. Rats were maintained at 90% of their free feeding weight throughout the experiments; food was given at the same time each day or after tests with drug or vehicle administration were completed on test days. Diet was a corn-based chow (Teklad Diet 2018; Harlan, Indianapolis, IN) and rats had free access to water except during test procedures. All procedures used in this study were approved by the Johns Hopkins Institutional Animal Care and Use Committee. The facilities adhered to the National Institutes of Health *Guide for the Care and Use of Laboratory Animals* and were AAALAC-approved.

### Drugs

(-)-trans-delta9-tetrahydrocannabinol (THC; 200 mg/ml in USP ethyl alcohol 95%) and cannabidiol (synthetic) were provided by the National Institute on Drug Abuse (NIDA) Drug Supply Program. THC and CBD were mixed with 100% sesame oil using sonication and vortex for an oral suspension. Sesame oil was used as it dissolves lipid-soluble cannabinoids for increased bioavailability and is utilized in pharmaceutical oral formulations of THC and CBD (e.g., Marinol and Epidiolex)(Zgair et al., 2016). THC (1, 3, 5.6, 10, and 20 mg/ml), CBD (3, 10, and 30 mg/ml), and the sesame oil vehicle were administered via oral gavage at a volume of 1 ml/kg. Morphine sulfate (Sigma Aldrich, St. Louis, MO, USA) was dissolved in 0.9% sterile saline for final doses of 5, 10, and 20 mg/ml, and was administered at 1 ml/kg via IP injection; morphine doses were calculated based on the salt.

### Study design

The effects of administration of THC or CBD via oral gavage (p.o.) were tested in male and female rats (N=24, 6 per sex/drug group). Separate groups of rats were used for THC and CBD tests. Treatments and behavioral tests were conducted once each week, with a minimum of 7 days between each THC/CBD dose. Vehicle treatments and tests were interspersed throughout the treatment period to assess carry-over drug effects and control for possible baseline shifts over time. THC injections were administered in a blinded, within subject Latin square design (0-20 mg/kg). CBD was administered in descending order. The total testing period was 9 and 5 weeks for THC and CBD, respectively. Behavioral testing occurred at baseline (pre-treatment) and at a series of time points for up to 7 hours following treatment with THC, CBD, or vehicle.

In a separate group of rats (N=8, 4 per sex), the effects of morphine (5-20 mg/kg) administered via IP injection were evaluated as a positive control for comparison of pain sensitivity outcomes. Morphine was administered in a blinded, within subject Latin square design (0-20 mg/kg) with testing occurring once per week, with a minimum 7 days apart. The total morphine testing period was 4 weeks.

### Cannabinoid Tetrad Test Battery

#### Antinociception (thermal pain sensitivity): Tail Flick Assay

Thermal pain sensitivity was assessed using the tail flick (TF) assay. In this test, the distal end of the rat’s tail (~50 mm from the tip) is exposed to radiant heat from a precise photobeam (Harvard Apparatus, Cambridge, MA, USA) and latency (s) to respond to the heat stimulus by flexion of the tail is recorded. The maximum duration was limited to 10s. Prior to testing, the radiant heat setting was calibrated to achieve a group average baseline latency of 5s. Baseline TF latencies were obtained immediately prior to drug administration and averaged across test weeks. Antinociception was calculated as percent of maximum possible effect (% MPE= [(test TF latency– baseline TF latency)/(maximum TF latency – baseline TF latency)] x 100). Rats were tested at 3 time points post oral administration: 60 min, 120 min, and 300 min for THC, CBD, and vehicle.

#### Antinociception (mechanical pain sensitivity: von Frey Test

The von Frey test was used to assess THC effects on mechanical pain sensitivity. Rats were placed in clear plastic cubicles on an elevated screen platform. Each hind paw is probed with von Frey filaments (9 filaments, 0.6-15.0 g, beginning with 2.0g) on the plantar surface for 3s. The presence or absence of a response (nocifensive hind paw flexion reflex) is recorded. If no response occurs, a stronger stimulus is presented otherwise the next weaker stimulus is applied. This up-down process is repeated 4 times after the first change in response, and the 50% threshold for paw withdrawal is determined by the individual response pattern and the force of the last von Frey filament tested (Chaplan et al., 1994; Dixon, 1991). Antinociception is demonstrated by an increase in the 50% mechanical withdrawal threshold. Baseline thresholds were obtained immediately prior to drug administration and averaged across test weeks. Antinociception was calculated as percent of maximum possible effect (% MPE= [(test threshold– baseline threshold)/(maximum threshold – baseline threshold)] x 100). Rats were tested at 3 time points post-drug administration, following the tail flick assay: 75 min, 135 min, and 315 min for THC, CBD, and vehicle.

### Locomotor Activity

Rats were placed in standard activity chambers (San Diego Instruments Inc.) where automated activity data was collected using a 4 x16 photobeam array. Tests were conducted in the dark with a white noise machine. Locomotor chambers interfaced with a computer running Photobeam Activity System (PAS) software that automatically recorded all beam interruptions, central peripheral activity, ambulation movements, fine movements, and time stamped (x,y) positions. Locomotor tests were 10-min in duration and occurred at 30, 150, and 270 min post oral administration of THC, CBD, or vehicle.

#### Catalepsy: Bar Test

Catalepsy, or immobilization, was tested using a bar test (Sanberg et al., 1988). Each individual chamber contained a bar apparatus 12 cm high and 5 mm in diameter (Med Associates, St. Albans, VT). Rats were placed with both front paws in contact with the bar.Experimenters blinded to the treatment condition timed the duration of contact with the bar, stopping the trial once both paws were removed. Total time (seconds) spent in contact with the bar, up to a maximum 180s was recorded. Trials were repeated after 2-min for a total of 3 trials per test. An average time immobile across the 3 trials was used as a measure of catalepsy. Catalepsy was measured at baseline, 90-min, and 330-min post oral administration of THC or vehicle.

### Body temperature

Body temperature was determined with a digital rectal thermometer with a lubricated flexible probe across 3-5 time points on the test day. Temperature was measured at 60, 120, 210, 300, and 420 min after THC or vehicle administration, or at 60, 120, 300 min after CBD or vehicle administration.

### Statistical Analysis

Thermal and mechanical pain sensitivities were calculated as percent of maximum possible effect (% MPE= [(test threshold-baseline threshold)/(maximum threshold – baseline threshold)] x 100). Maximum thresholds for the tail flick and von Frey assays were 10 and 15, respectively. An average of baseline tests were used in these calculations. Locomotor activity was calculated as a percent change from each animal’s average of weekly vehicle tests. Time immobile (s) in the catalepsy test was logarithmically transformed (ln) prior to analysis (Ferre et al., 1990). Outcome measures analyzed included: %MPE, body temperature change (°C), locomotor activity distance traveled (% of vehicle), and time immobile (ln(s)) in the catalepsy test. Three-way repeated measures ANOVAs were conducted with sex as a between subjects variable and dose and time as within subject variables; for interpretation of interactions with time, Bonferroni post-hoc tests were used. Follow up two-way ANOVAs (sex as a between subjects variable and dose as a within subject variable) were conducted within each time point. Dunnett’s post hoc tests were used to analyze differences in outcomes between THC or CBD dose/condition and vehicle. In the event of a main effect or interaction with sex, post-hoc comparisons between males and females were determined with Sidak’s test. Post-hoc results with sexes collapsed are reported in the text, figures show the average data as well as points for males and females separately. ED50 values were calculated with nonlinear regression using data from the time point where maximum effects were observed. All statistics were performed with Statistica 11 and Graphpad Prism 9.

## Results

### Effects of oral THC

#### Thermal Pain Sensitivity

Oral THC administration produced antinociception in the tail flick assay (**Fig. 1A**) as confirmed by a significant main effect of THC dose (F(5, 45)= 6.67, p<0.001). There was also a main effect of time (F(2, 18) =8.13, p<0.01), as %MPE increased across the testing period, and an interaction of time × sex (F(2, 18) =4.15, p<0.05). The average %MPE of male rats increased over the time course (p’s<0.05) while female %MPE was equivalent across the 3 tests. When analyzed within each time point, THC (5.6-20 mg/kg) produced thermal antinociceptive effects at 120-min (F(3.47, 40.91)= 4.27, p<0.01) and 300 min (F(2.46, 24.07)= 6.11, p<0.01).

**Figure 1.**
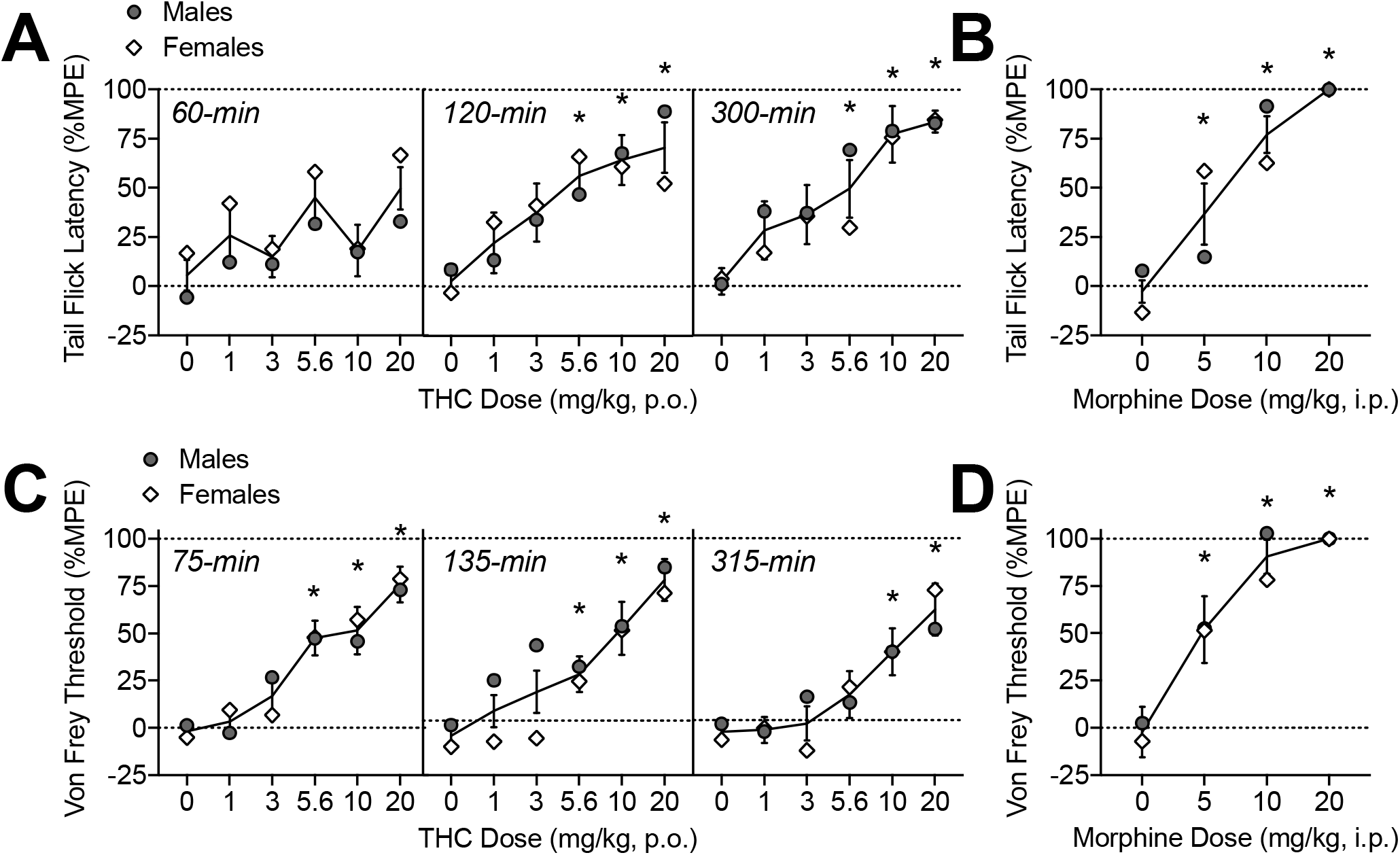
Oral THC effects on thermal (A) and mechanical (C) pain sensitivity. Morphine (i.p.) comparison (B, D). The time of data collection after oral administration is noted in *italics*. Asterisks (*) represent a significant difference from vehicle. Data are Mean ±SEM; N=6/sex for THC, N=4/sex for morphine.

#### Mechanical Pain Sensitivity

Oral THC administration produced antinociception in the von Frey test (**Fig. 1B**) as confirmed by a significant main effect of THC dose (F(5, 50) =3.39, p<0.05). There was also a main effect of time (F(2, 20) =11.43, p<0.001), an interaction of time × sex (F(2, 20) =4.81, p<0.05), and an interaction of THC dose × sex (F(10, 100) =4.51, p<0.001). Von Frey thresholds (%MPE) overall were declining by 300-min compared to the first test (p<0.05), particularly in females, indicating effects returning to baseline. When analyzed within each time point, THC (5.6-20 mg/kg) produced significant mechanical antinociceptive effects at 75-min, (F(3.67, 36.69)= 12.51, p<0.001) and 135-min post-administration (F(3.19, 31.90)= 10.54, p<0.001). At 315-min, the highest doses of THC (10-20 mg/kg) continued to show mechanical antinociceptive effects (F(3.45, 34.49)= 8.153, p<0.001).

#### Body Temperature

Oral THC administration reduced body temperature dose-dependently (**Fig. 2**) as confirmed by a main effect of THC dose (F(5, 50)= 10.84, p<0.001), time (F(4, 40)= 13.40, p<0.001), and an interaction of THC dose × time (F(20, 200)= 2.44, p<0.001). When analyzed within each time point, oral THC decreased body temperature at 120-min (20 mg/kg), 210-min (10-20 mg/kg), 300 (1, 5.6-20 mg/kg), and 420-min (3, 10-20 mg/kg) post-administration (F’s between 4.18-11.43, p’s<0.05). There were main effects of sex at 120, 210, and 300 (F’s (1,10) = between 9.93-14.63); under vehicle conditions, males showed slightly increasing temperatures (+0.14-0.19°C) throughout the course of the testing period compared to females (−0.04-0.06°C; p’s<0.05). At 420-min post oral THC, there was an interaction of THC dose and sex (F(5, 50)= 2.87, p<0.05), though post-hoc tests did not isolate sex differences to any specific THC dose.

**Figure 2.**
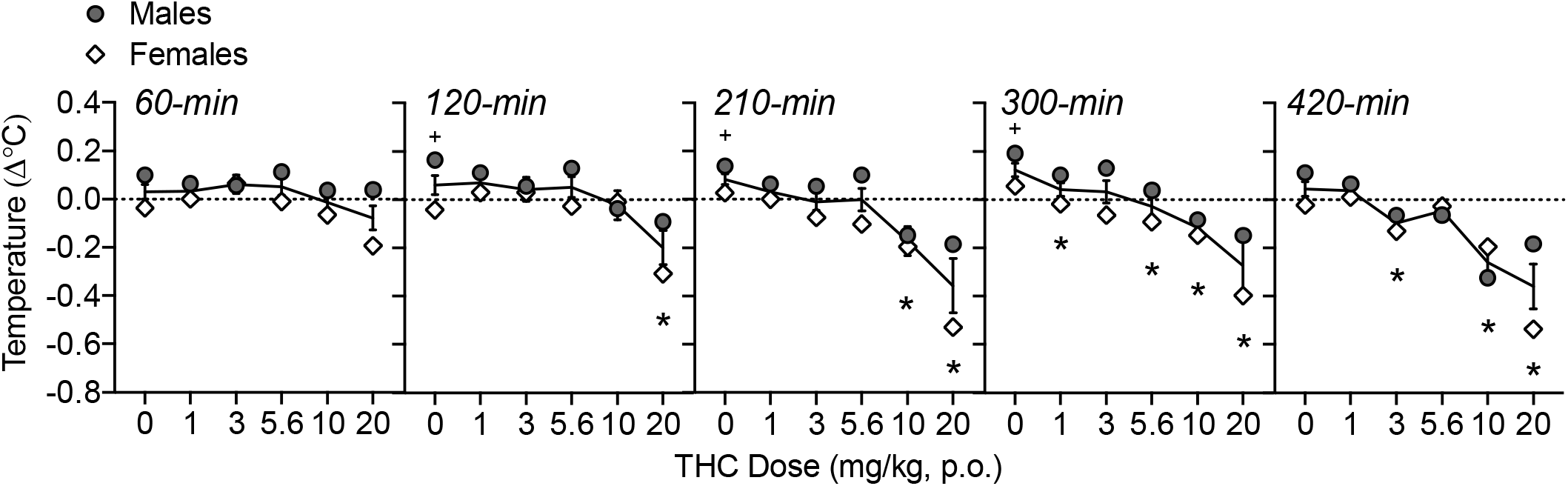
Oral THC effects on body temperature change from baseline (Δ°C). The time of data collection after oral administration is noted in *italics*. Asterisks (*) represent a significant difference from vehicle. Plus sign (+) indicates a sex difference. Data are Mean ±SEM; N=6/sex.

#### Locomotor Activity

THC modulated locomotor activity over the testing period (**Fig. 3**) as confirmed by a main effect of THC dose (F(5, 50)= 3.38, p<0.05) and time (F(2,20)= 11.81, p<0.001). There were also interactions of THC dose × time (F(10, 100)= 4.60, p<0.001) and time × sex (F(2, 20)= 4.54, p<0.05). When collapsed across dose to assess the time × sex interaction, females’ activity was lower at the last time point (270-min) compared to the first time point (30-min; p<0.05). At 30-min post oral administration, THC increased locomotor activity at 5.6 and 20 mg/kg (F(3.29, 32.93)= 6.61, p<0.01). There was an effect of sex at 30-min (F (1, 10)= 5.77, p<0.05), with females showing higher activity than males after 10 mg/kg THC (p<0.05). At 150-min post oral THC administration, there was a main effect of THC dose on locomotor activity (F(3.01, 30.13)= 2.86, p=0.05), though post-hoc tests did not indicate any specific doses were different than vehicle. By 270-min, THC reduced locomotor activity at 3 and 5.6 mg/kg (F(3.47, 34.74)= 3.57, p<0.05).

**Figure 3.**
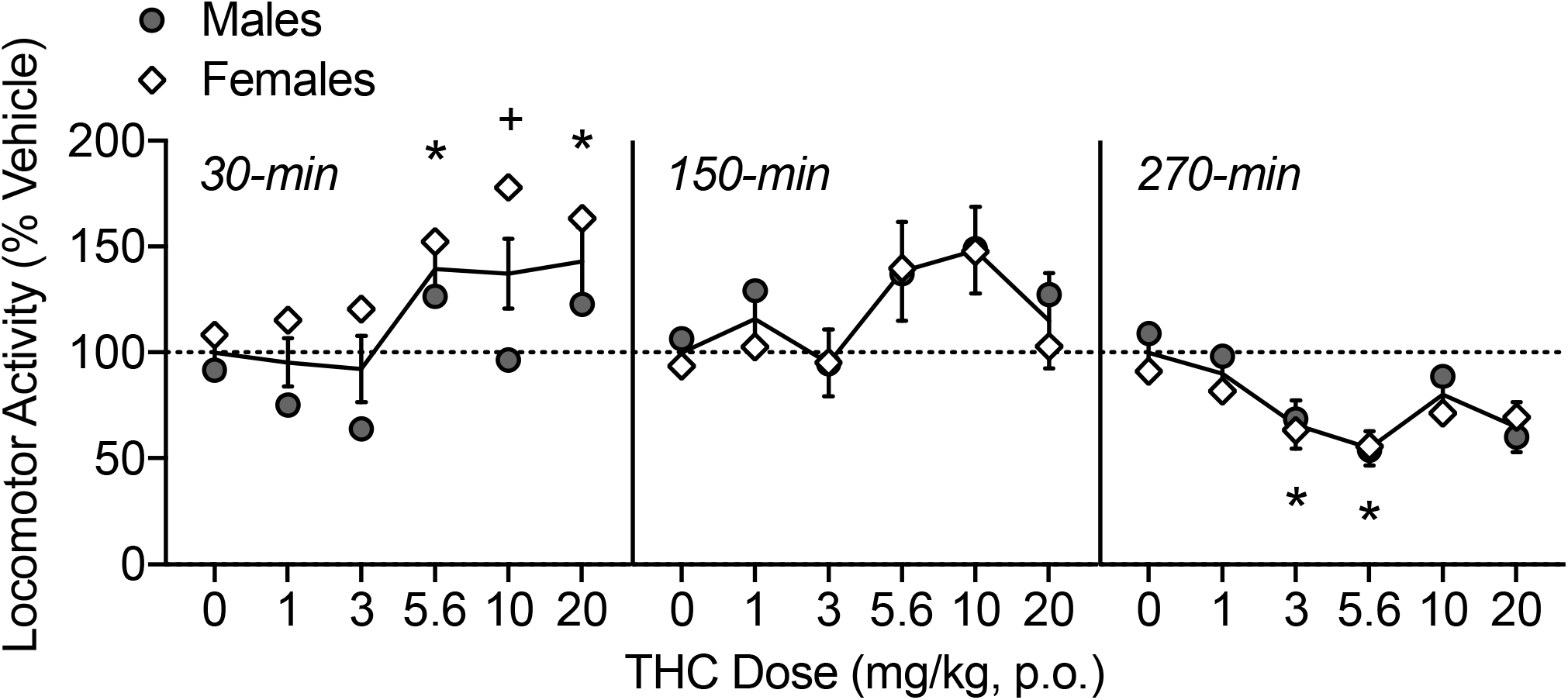
Oral THC effects on locomotor activity, shown as percent of vehicle distance traveled. The time of data collection after oral administration is noted in *italics*. Asterisks (*) represent a significant difference from vehicle. Plus sign (+) indicates a sex difference. Data are Mean ±SEM; N=6/sex.

#### Catalepsy

High doses of THC caused increases in catalepsy over the testing period (**Fig. 4**) as confirmed by a main effect of THC dose (F(5, 50)= 13.54, p<0.001) and time (F(1, 10)= 5.55, p<0.05). At 90-min, 20 mg/kg THC increased catalepsy (F(3.26, 32.57)= 11.20, p<0.001), and at 330-min, 10-20 mg/kg THC increased catalepsy (F(3.04, 30.40)= 6.68, p<0.001).

**Figure 4.**
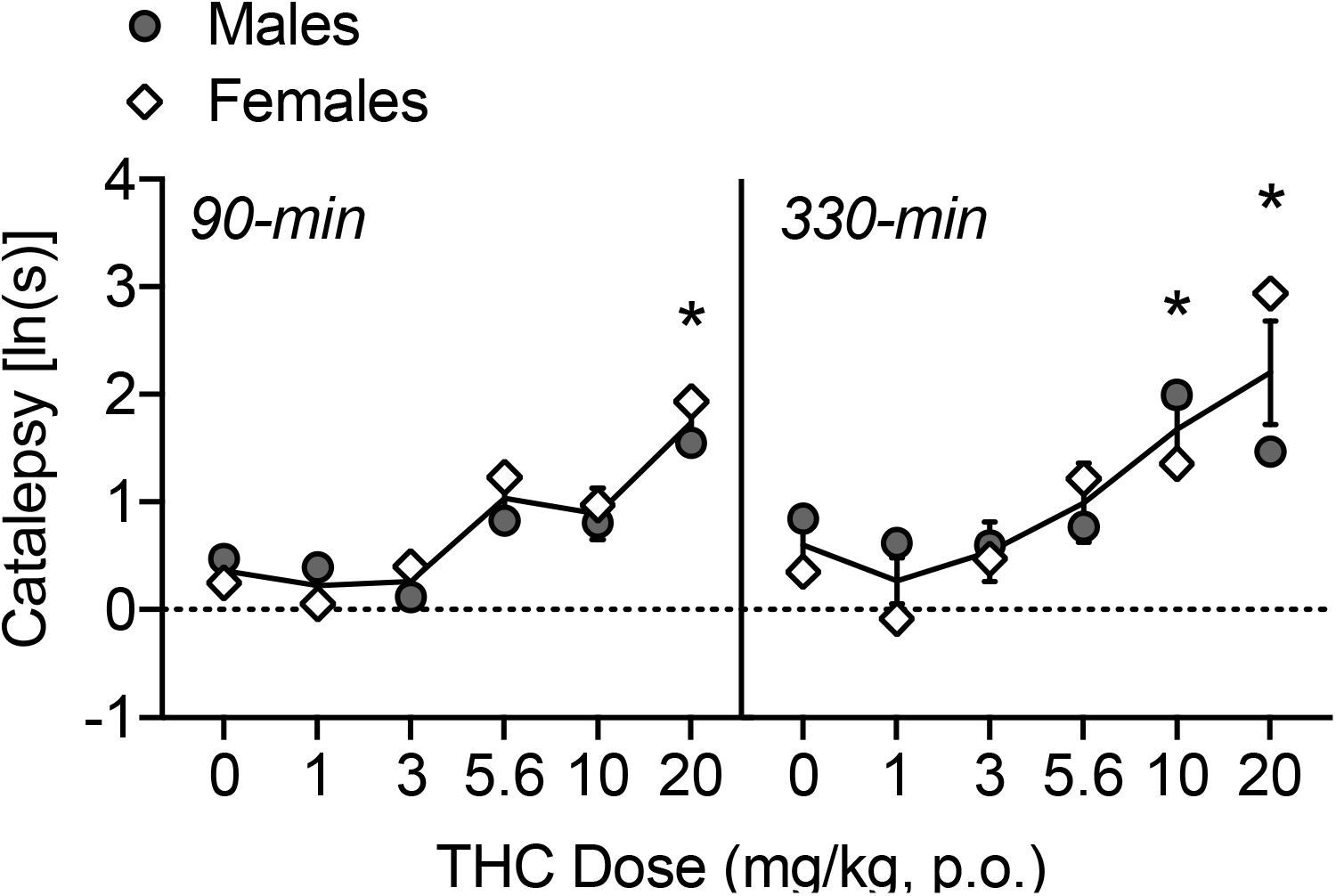
Oral THC effects on catalepsy. The time of data collection after oral administration is noted in *italics*. Asterisks (*) represent a significant difference from vehicle (males or females). Data are Mean ±SEM; N=6/sex.

#### ED50 values

ED50 values were calculated at the time point where effects were maximal (see **Table 1**). The ED50 values for THC’s thermal antinociceptive effects were 1.66 and 4.34 mg/kg for males and females, respectively. In comparison, the ED50 values observed for morphine in the tail flick test were 9.00 and 4.76 mg/kg for males and females. For mechanical antinociceptive effects, ED50 values were 4.32 and 6.40 mg/kg for males and females, which was less potent than morphine (ED50 = 2.42 and 2.96 mg/kg for males and females). Notably, oral THC was less efficacious at producing thermal and mechanical antinociception, with maximum responses in the tail flick and von Frey test around 80% compared with 100% efficacy observed with morphine.

**Table 1.**
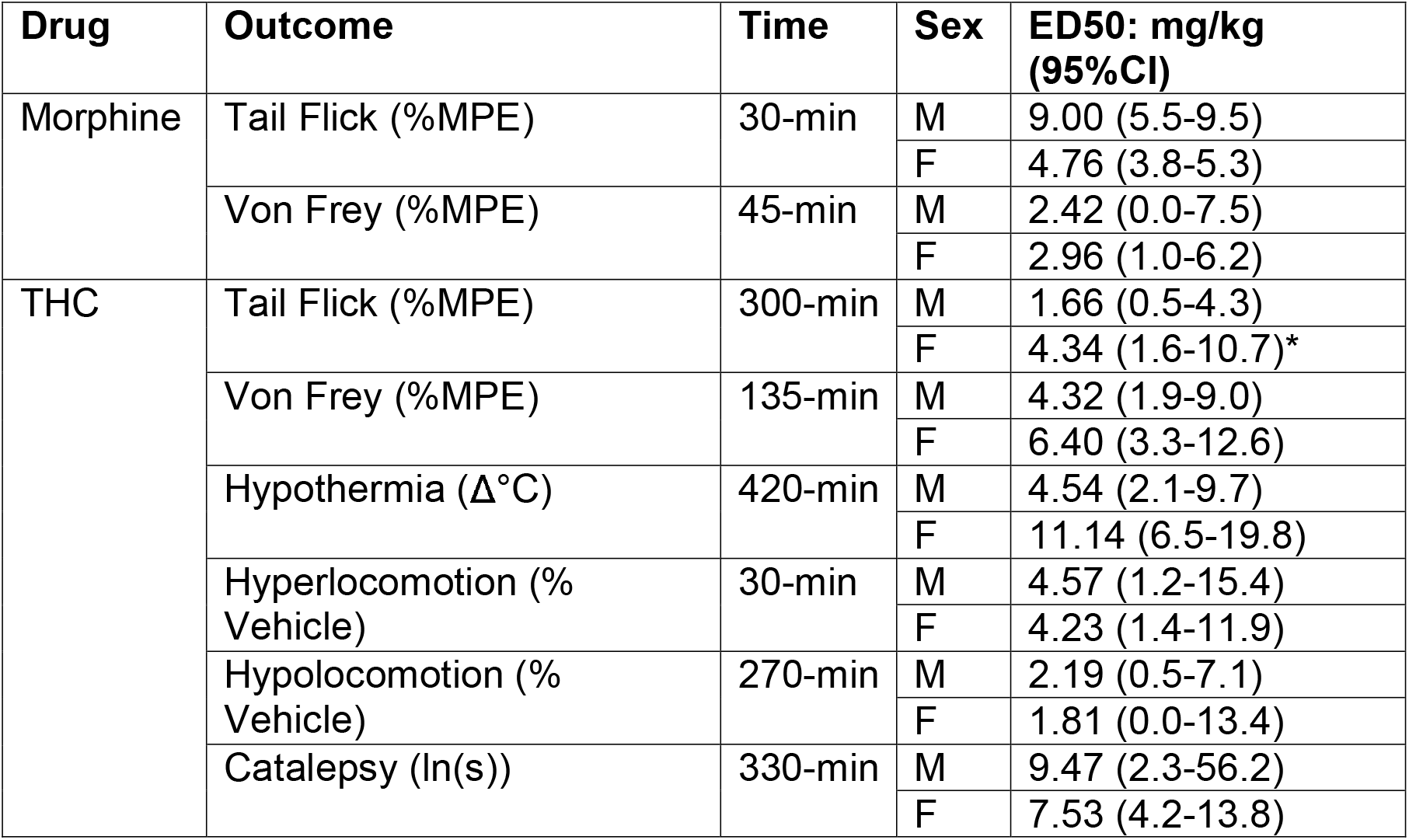
ED50 values of THC and Morphine. ED50 was calculated for the time point in which maximal effects were observed. An asterisk (*) is used to denote non-overlapping 95% confidence intervals between males and females.

The IC50 for hypothermic effects of THC were calculated to be 4.54 and 11.14 mg/kg THC for males and females, respectively. IC50 values for hyperlocomotive effects (observed at 30-min) were 4.57 and 4.23 mg/kg for males and females, while the ED50 values for hypolocomotive effects (270-min) were 2.19 and 1.81 mg/kg for males and females. ED50 for cataleptic effects was 9.47 and 7.53 mg/kg for males and females, respectively.

### Effects of oral CBD

#### Thermal and Mechanical Pain Sensitivity

Oral CBD administration had modest effects on pain sensitivity as measured in the tail flick assay and von Frey test (**Fig. 5A-B**). There was a main effect of CBD dose on tail flick latencies (%MPE) (F(3, 30)= 3.73, p<0.05); however, post-hoc tests did not isolate any doses that had significant differences from the vehicle condition at any time point tested. There was a CBD dose × time × sex interaction on von Frey thresholds (%MPE; F(6, 60)= 2.87, p<0.05). CBD (30 mg/kg) increased pain sensitivity at 315-min (F(1.88, 18.75)= 4.29, p<0.05). At 315-min, there was also an interaction of CBD dose × sex (F(3, 30)= 3.13, p<0.05), and while there were no significant sex differences determined by post-hoc tests, this interaction was likely driven by a greater effect of 30 mg/kg CBD in increasing mechanical nociception in females at this time point.

**Figure 5.**
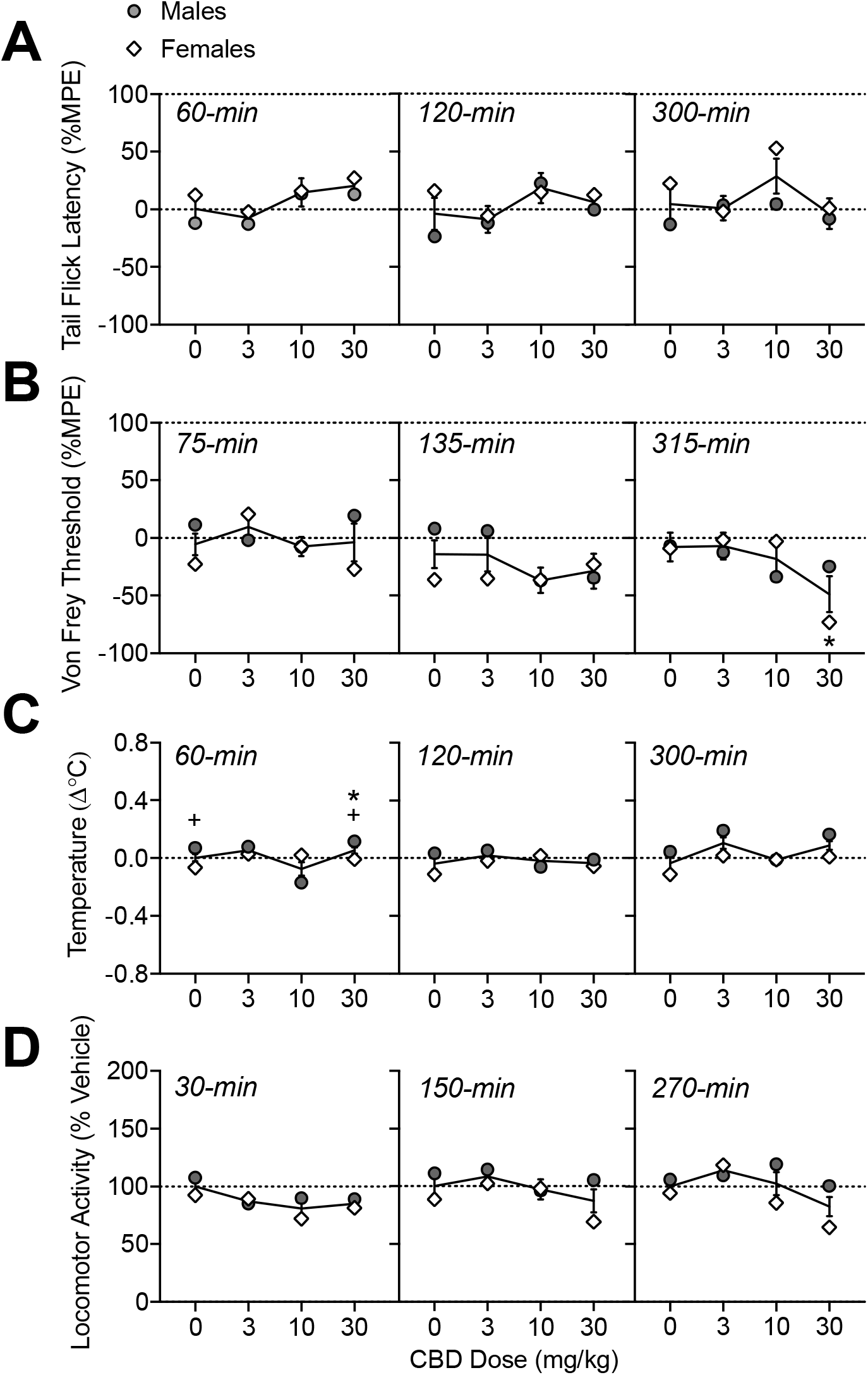
Oral CBD effects on thermal (A) and mechanical (B) pain sensitivity, body temperature (C), and locomotor activity (D). The time of data collection after oral administration is noted in *italics*. Asterisks (*) represent a significant difference from vehicle. Plus sign (+) indicates a sex difference. Data are Mean ±SEM; N=6/sex.

#### Body Temperature

Oral CBD administration altered temperature in a sex-dependent manner (**Fig. 5C**; main effects of CBD dose F(3, 30)= 6.31, p<0.001; sex: F(1, 10)= 9.43, p<0.05; and a CBD dose × sex interaction: (F(3, 30)= 8.50, p<0.001)). At 60-min, 30 mg/kg CBD increased body temperature (F(1.33, 13.33)= 6.73, p<0.05). There was also an interaction of CBD dose × sex (F(3, 30)= 10.48, p<0.001), with males showing higher temperature changes than females under vehicle conditions and after 30 mg/kg CBD, though these differences were small (mean difference: 0.12-0.14°C). At 300-min post oral CBD administration, there was a main effect of CBD dose (F(1.72, 17.15)= 3.79, p<0.01), though post-hoc tests did not indicate differences from vehicle for any specific dose.

#### Locomotor Activity

Oral CBD administration modulated locomotor activity (**Fig. 5D**; main effect of CBD dose: (F(3,30)= 9.59, p<0.001) and interactions of CBD dose × sex (F(3, 30)= 5.00, p<0.001), CBD dose × time (F(6, 60)= 2.75, p<0.05), and CBD dose × time × sex (F(6, 60)= 3.18, p<0.01). Analysis within each time point determined that CBD (10-30 mg/kg) reduced locomotor activity at 30-min (F(2.60, 25.99)= 6.89, p<0.01. At 270-min post oral CBD administration, there was a main effect of CBD dose (F(2.28, 22.83)= 5.72, p<0.01); though post-hoc tests did not indicate differences from vehicle for any specific dose. There was also a CBD dose × sex interaction (F(3, 30)= 3.75, p<0.05); likely due to lower activity in females than males at 10-30 mg/kg, though sex differences were not significant according to posthoc tests.

## Discussion

Oral THC produced typical behavioral effects in the classic tetrad: antinociception, hypothermia, hypolocomotion, and catalepsy. These outcomes were mostly equivalent in males and females and time course of effects were quite prolonged, lasting up to 7 hours post administration.

Dose-dependent antinociceptive effects in the tail flick test were observed at 120 and 300 minutes following oral THC administration, with maximal effects seen at 300-min. There were sex differences in the time course of antinociception: in males, antinociceptive effects of THC increased across the testing period, while antinociception in females was equal across time points. The potency of the thermal antinociceptive effects at 300-min, when effects were maximal, was higher in males compared to females. Dose-dependent antinociceptive effects were also observed in the von Frey test when tested at 75, 135, and 315 minutes following oral THC administration, with maximal effects seen at 135-min. There were sex differences in the time course of mechanical antinociception: in females, antinociceptive effects were lower after 315-min compared to the 75-min time point, indicating a return to baseline levels, while the %MPE in males was equal across time points. Taken together, antinociceptive effects were observed in both tests of pain sensitivity, though overall, mechanical antinociceptive effects were observed earlier and reached their maximum at an earlier time point compared with thermal antinociceptive effects, which peaked later. However, this time course of antinociceptive effects was sex-dependent: males primarily showed a slower onset of thermal antinociceptive effects and prolonged mechanical antinociception compared with females, who had a faster onset (thermal antinociception) and offset (mechanical antinociception).

Notably, when compared with morphine (20 mg/kg), oral THC (20/mg/kg) was less efficacious at producing thermal and mechanical antinociception, with maximum responses in the tail flick and von Frey test around 80% compared with 100% efficacy observed with morphine. The potency of oral THC and IP morphine to produce thermal antinociception were roughly equivalent in females (ED50s ~ 4 mg/kg), while the ED50 in males was lower for oral THC compared with morphine (1.66 vs 9.00 mg/kg, respectively). The potency of morphine was around two-fold higher than THC for producing mechanical antinociceptive effects (ED50 ~2 and 3 mg/kg for morphine and ~4 and 6 mg/kg for THC, in males and females respectively).

In the current study, there were modest sex differences observed in the antinociceptive effects of oral THC, particularly in the time course of effects. In studies using IP injected THC, we and others have previously shown sex differences in antinociception, where females show greater sensitivity to IP THC (i.e. effects at lower doses, greater magnitude of effects, longer lasting effects) compared to males (Craft et al., 2019; Moore et al., 2021; Tseng and Craft, 2001). In contrast, vapor exposure to THC produced equivalent antinociceptive effects in male and female rats (Moore et al., 2021). Our current results indicate a greater potency of oral THC to produce thermal antinociceptive effects in males compared with females. In a separate study using oral THC, sex differences in antinociceptive effects were observed in a neuropathic pain model, where chronic oral THC reduced hypersensitivity in male rats, but had no effect in female rats (Linher-Melville et al., 2020). In a study of volitional consumption of oral THC in gelatin (~2 mg/kg), equivalent antinociception was observed in male and female rats when tested once immediately after 1-hr access to THC (Kruse et al., 2019). Results from the current study demonstrate that the magnitude and/or expression of sex effects may depend on the time course of drug effects when using oral administration.

We observed modest, orderly decreases in body temperature over time, with the temperature nadir of −0.4°C observed 7 hours after oral THC administration. In comparison, in our recent study (Moore et al., 2020), we observed temperature nadirs of −1.0°C, 300-min after administration of 20 mg/kg IP THC, and and −0.7°C, 60-min after ~100 mg THC vapor exposure. A separate study measuring body temperatures after voluntary oral consumption of THC in gelatin (~2 mg/kg) showed a similar magnitude of decrease when measured immediately following 1-hr access in adolescent male and female rats (Kruse et al., 2019).

We observed both hyper- and hypolocomotor effects of oral THC that depending on the time of the activity test post THC administration. When compared to vehicle, oral administration of THC increased locomotor activity at 30 min followed by reductions in locomotor activity (hypolocomotion) after 5 hours. In mice, IP injected THC produces hypolocomotion, observable immediately and lasting up to 420-min post administration (Martin et al., 1991; Metna-Laurent et al., 2017; Puighermanal et al., 2013; Tai et al., 2015; Wiley et al., 2021). In contrast, both hypolocomotive and hyperlocomotive effects of THC have been reported in rats. A study using IP THC observed “triphasic” effects, where very low doses and high doses reduced locomotor and mid-range doses stimulated activity (Sanudo-Pena et al., 2000). Studies utilizing THC vapor exposure have observed either no effect (Javadi-Paydar et al., 2018; Nguyen et al., 2018) or hypolocomotion (Nguyen et al., 2016) when measuring home-cage activity for 3 hours after vaporized THC exposure. In a study of voluntary oral consumption of low dose THC (~2 mg/kg) in gelatin, hyperlocomotion was observed immediately following 1-hr access to THC in a 10-min activity test, and increased activity was similar in males and females (Kruse et al., 2019). A study of male and female adolescent rats given low doses (1 and 5 mg/kg) of oral THC in sesame oil found no effects on locomotor activity when tested from 40-100 min post administration (Dow-Edwards and Zhao, 2008). In a separate study using rats, 10 mg/kg oral THC in sesame oil overall reduced locomotor activity in a 5-min test when measured 120 minutes after administration (Hlozek et al., 2017). Our results also indicate sex differences in locomotor effects of THC, where hyperlocomotion at 30-min was higher in females. A previous study found that subcutaneously injected THC (30 mg/kg) produced hypolocomotion in male, but not female rats, when tested 30-min after injection (Marusich et al., 2014); though other studies using IP injection have observed no sex differences when tested after 30-min (Wiley et al., 2007). In a study of vaporized THC, hypolocomotion was observed in male, but not female rats (Javadi-Paydar et al., 2018). To our knowledge, this is the first study reporting both increases and decreases in activity after oral THC administration, and the first to observe sex differences in locomotor effects of oral THC.

In the current study, catalepsy was observed at the highest doses of oral THC tested (10-20 mg/kg) and this peaked at 5.5h after administration. Prior studies have shown that cataleptic effects of THC depend on route of administration (Marshell et al., 2014). While IP THC and synthetic CB1 agonists reliably produce catalepsy, cataleptic effects were not observed in male mice after exposure to vaporized THC or synthetic cannabinoids JWH-018 and JWH-073 (Marshell et al., 2014). The present data concurs with what has been shown using IP injection: catalepsy produced by 10 mg/kg IP THC was shown to last up to 6 hours in male rats (Prescott et al., 1992). A study using oral THC (50 mg/kg) observed catalepsy in female mice when tested at 60-min post administration (Burstein et al., 1987). Sex differences in the cataleptic effects of THC have also been reported: 10 mg/kg IP THC produced greater amounts of catalepsy in female rats compared to male rats, and effects peaked at 60-min post injection (Tseng and Craft, 2001), likely due in part to sex differences in THC metabolism (Tseng et al., 2004). In the present study, there were no sex differences observed in the cataleptic effects of oral THC.

In the present study, oral CBD had only modest effects on behaviors in the tetrad, which was expected. No antinociceptive effects of CBD were detected in the tail flick test, and in fact, increased pain sensitivity was observed after the highest dose of CBD (30 mg/kg) in the von Frey test 300-min after administration. In the von Frey assay, CBD effects were not dose-dependent or consistent across time points. While CBD has been shown to reduce or prevent hyperalgesia in inflammatory and neuropathic pain models (King et al., 2017; Mlost et al., 2020), acute antinociceptive effects were not observed in normal animals after CBD when given IP (1.25-30 mg/kg) (Britch et al., 2017) or after vaporized exposure (Javadi-Paydar et al., 2019). To our knowledge, oral CBD effects on acute pain sensitivity has not been assessed in rats; replication of these effects is necessary for determining whether oral CBD reliably increases mechanical nociception in female rats.

We observed effects on temperature in a sex-dependent manner; however, these changes in temperature were small (<0.1°C) and would not be considered meaningful. CBD injected IP has been shown to have modest hypothermic (Long et al., 2010) or no effects (Varvel et al., 2006) on body temperature. Vaporized CBD has been shown to reduce body temperature in rats (Javadi-Paydar et al., 2019; Javadi-Paydar et al., 2018). A lack of hypothermic effects in the present study suggests that reductions in body temperature often observed after IP or vaporized CBD is less likely to be observed under conditions of oral CBD administration.

We observed effects of CBD on locomotor activity: specifically, hypolocomotion 30-min following oral administration of 10-30 mg/kg CBD. Locomotor activity after 270-min was modulated by CBD in a sex-dependent manner, where female’s activity was overall lower compared to males, though sex effects were not isolated to any particular dose. Similar to THC, effects of CBD on locomotor activity seem to depend on dose and perhaps route of administration, with studies in mice reporting no effects at low doses (1-5 mg/kg), increases and decreases in locomotor activity after high CBD doses (10-30 mg/kg) via injection (IP, intravenous) (Long et al., 2010; Varvel et al., 2006). In rats administered CBD via IP injection, hyperlocomotive effects were observed after 10 and 30 mg/kg, but only after 240-360 min post administration, and these effects were equivalent in males and females (Britch et al., 2017). Vaporized CBD has been shown to produce hypolocomotion immediately following 1-hr exposure, which returned to control levels after 30-min (Javadi-Paydar et al., 2019). A study in rats did not observe any effects of 10 mg/kg oral CBD when tested once, 120-min after administration (Hlozek et al., 2017). Taken together with results from the present study, route of administration may have differential effects on locomotor response to CBD and the time course of those effects.

## Conclusions

The findings from the current study on the time course of dose-effects of orally administered cannabinoids in the tetrad provide foundational data for future studies for selection of THC and CBD dose and timing of behavioral measures when using the oral route of administration. Results from this study, in conjunction with our prior study assessing time course of behavioral effects of IP and vaporized THC (Moore et al., 2020), demonstrate the importance of consideration of time course when evaluating behavioral effects, particularly when using a nontraditional route of drug administration.

## Acknowledgements

All experiments were supported by the National Institute on Drug Abuse of the National Institutes of Health grant numbers R21DA046154 (EW) and the Johns Hopkins University Dalio Fund in Decision Making and the Neuroscience of Motivated Behaviors (EW). The authors have no conflicts of interest to disclose.

## Abbreviations

(THC): Δ9-tetrahydrocannabinol
(CBD): Cannabidiol
(CB1R): cannabinoid receptor 1
(IP): intraperitoneal
(SC): subcutaneous
(IV): Intravenous
(TF): tail flick
(MPE): Maximum possible effect

